## Supplemental Materials for "Multivariate genome-wide association study on tissue-sensitive diffusion metrics identifies key molecular pathways for axonal growth, synaptogenesis, and astrocyte-mediated neuroinflammation"

Validated loci as druggable targets

To investigate how druggable our validated loci are, we queried the DGIdb ^45^ with all genes that mapped onto the validated loci (Supplementary Table 10-12). Among the validated loci, 218 of the N0 loci (64%) are druggable with 207 of them have known pharmacological interactions. This high degree of druggable targets was also 2observed among ND loci (185 loci are druggable and 175 of them have known interactions) and NF loci (125 loci are druggable and 125 of them have known interactions). The gene set analyses ^29^ show those druggable genes were highly enriched in brain tissues, especially hippocampus (*P*_bon_ = 8.0e-15, 1.5e-13, 2.5e-7 for N0, ND, and NF, respectively ) and amygdala (*P*_bon_= 1.4e-13, 6.2e-15, 2.5e-19 for N0, ND, and NF, respectively). Those druggable genes were evidently overlaps with genes identified in neuropsychiatric disorders and immune disorders, such as schizophrenia (*P*_bon_= 1.4e-8, 2.8e-9, 3.4e-6 for N0, ND, and NF), bipolar disorders (*P*_bon_= 9.3e-9, 4.3e-9, 1.2e-4 for N0, ND, and NF), and inflammatory bowel disease (*P*_bon_= 9.3e-14, 6.6e-14, 4.0e-12 for N0, ND, and NF).

*Average heritability across three RSI metrics*

Although each phenotypic PC can have heritabilities as high as 0.32 to 0.36 (Supplementary Figure 11), the average heritability of the combined PC for each imaging feature is modest. As the optimal heritability measures for high-dimensional multivariate measurements would be the mean heritability across PCs ^23^ (See Method), we found that the mean signal for N0, ND, and NF are 0.09 (95%CI 0.04 - 0.13), 0.06 (95%CI 0.02 - 0.10), and 0.05 (95%CI 0.01 - 0.09). Given the sum of the variance explained by the independent loci found to be significant, the discovered loci reached 59%, 60%, and 58% of the average SNP-heritabilities for N0, ND, and NF.

**Supplementary Table Legends**

**Supplementary Table 1. Characteristics of discovery and validation samples**

**Supplementary Table 2. Unique loci dicovered in current study, indexed based on the genomic physical positions.**

**Supplementary Table 3. Summary statistics, including mapped genes and overlaps with previous imaging GWAS, of unique loci for N0.**

**Supplementary Table 4. Summary statistics, including mapped genes and overlaps with previous imaging GWAS, of unique loci for ND.**

**Supplementary Table 5. Summary statistics, including mapped genes and overlaps with previous imaging GWAS, of unique loci for NF.**

**Supplementary Table 6. Top five enriched anatomical regions and their corresponding enrichment scores for N0 loci.**

**Supplementary Table 7. Top five enriched anatomical regions and their corresponding enrichment scores for ND loci.**

**Supplementary Table 8. Top five enriched anatomical regions and their corresponding enrichment scores for NF loci.**

**Supplementary Table 9. Region of Interests used in the regional enrichment analysis.**

**Supplementary Table 10. Druggable gene targets of N0 Locus**

**Supplementary Table 11. Druggable gene targets of ND Locus**

**Supplementary Table 12. Druggable gene targets of NF Locus**

**Figure Legends**

**Supplementary Figure 1. Histogram of the registration quality metrics across all imaging samples.**

**Supplementary Figure 2. Spatial distribution of N0.** The N0 metrics were averaged across discovery samples of UKB. The coronal sections were rendered slice by slice for every 8 mm. Upper left, first slice from the posterior region. Lower left, last slice from the frontal region.

**Supplementary Figure 3. Spatial distribution of ND.** The ND metrics were averaged across discovery samples of UKB. The coronal sections were rendered slice by slice for every 8 mm. Upper left, first slice from the posterior region. Lower left, last slice from the frontal region.

**Supplementary Figure 4. Spatial distribution of NF.** The NF metrics were averaged across discovery samples of UKB. The coronal sections were rendered slice by slice for every 8 mm. Upper left, first slice from the posterior region. Lower left, last slice from the frontal region.

**Supplementary Figure 5. Joint distribution of all three tissue sensitive diffusion metrics, N0, ND, and NF.**

**Supplementary Figure 6. Eigenvalues of principal components across three tissue sensitive diffusion metrics**

**Supplementary Figure 7. Eigenvectors of first 64 principal components of N0**. In each frame, the mid-coronal section of the brain was shown, starting from first PC (upper left) to the 64th PC (lower right), ordered accordingly.

**Supplementary Figure 8. Eigenvectors of first 64 principal components of ND**. In each frame, the mid-coronal section of the brain was shown, starting from first PC (upper left) to the 64th PC (lower right), ordered accordingly.

**Supplementary Figure 9. Eigenvectors of first 64 principal components of NF**. In each frame, the mid-coronal section of the brain was shown, starting from first PC (upper left) to the 64th PC (lower right), ordered accordingly.

**Supplementary Figure 10. Genome-wide signal overlaps between tissue sensitive diffusion metrics and other GWAS, including immune disorders ^65^, schizophrenia ^62^, attention deficit hyperactivity disorder ^66^, bipolar disorder ^60^, cross-psychiatric-disorders ^58^, Alzheimer's disease ^64^, educational attainment ^52^, and risk-related behaviors ^55^**

**Supplementary Figure 11. SNP-heritability of each principal component across three tissue sensitive diffusion metrics, estimated with LD score regression.**

Supplementary Figure 1

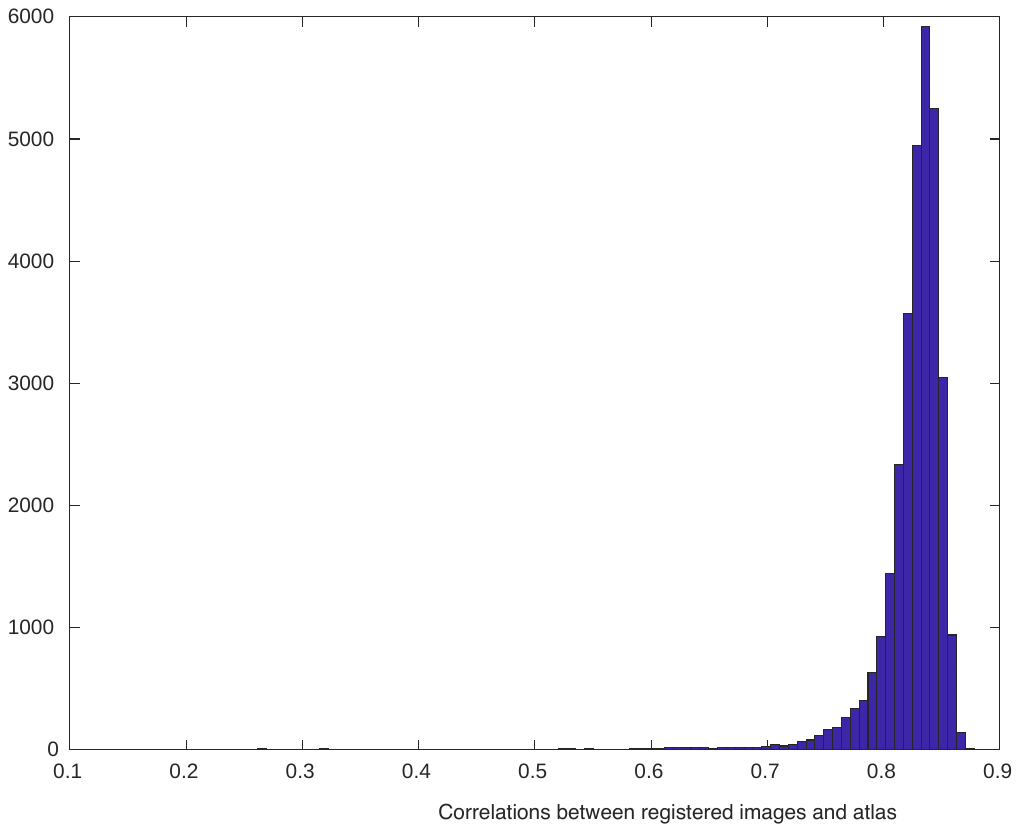

Supplemental Figure 2

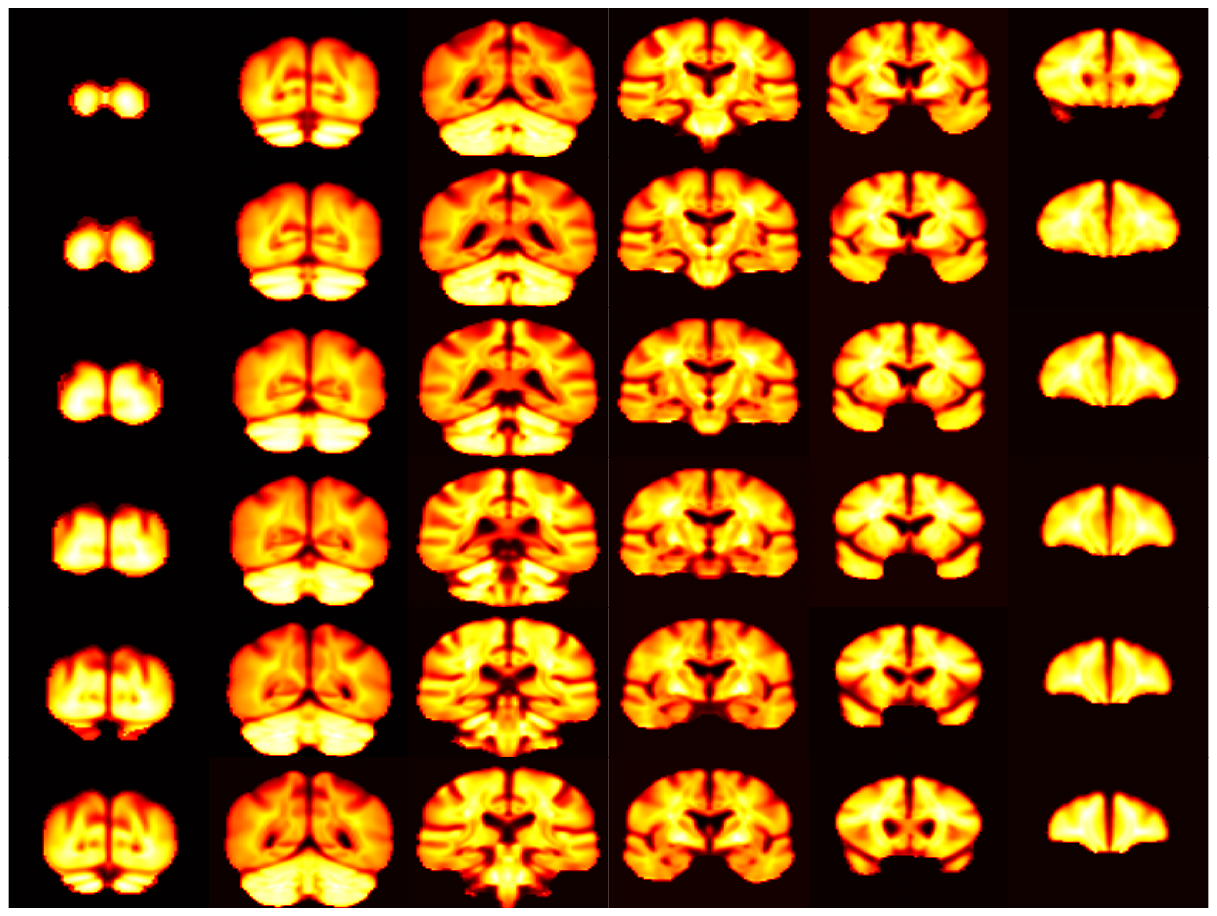

Supplemental Figure 3

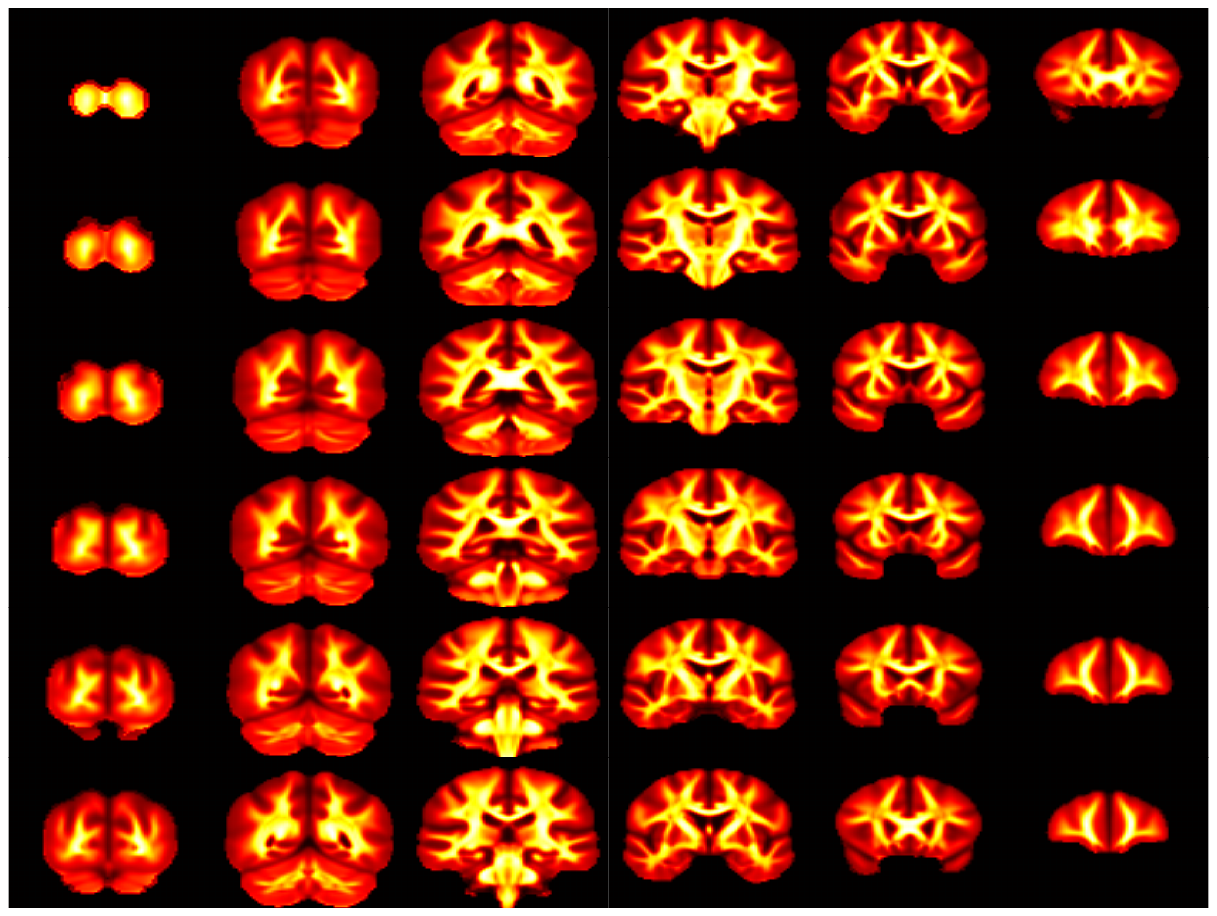

Supplemental Figure 4

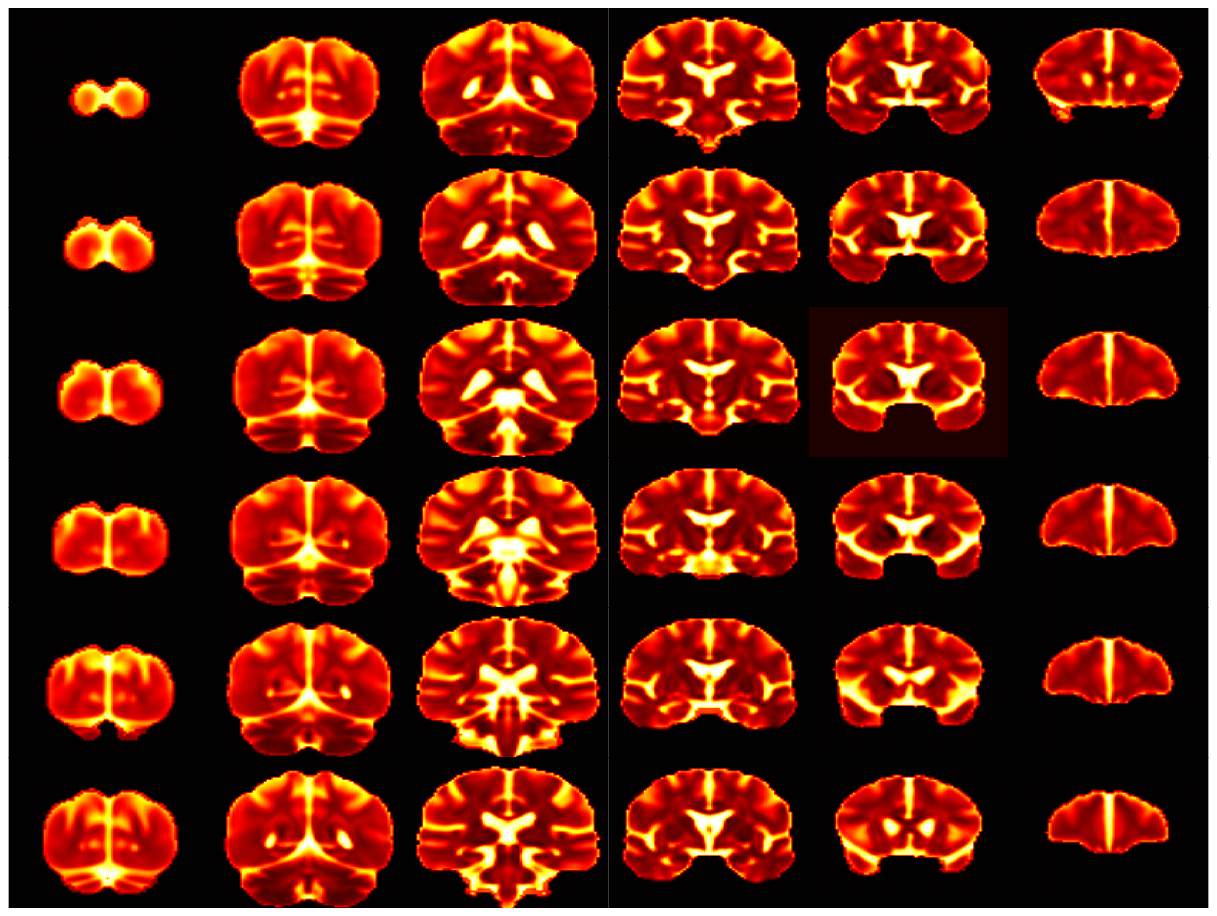

Supplemental Figure 5

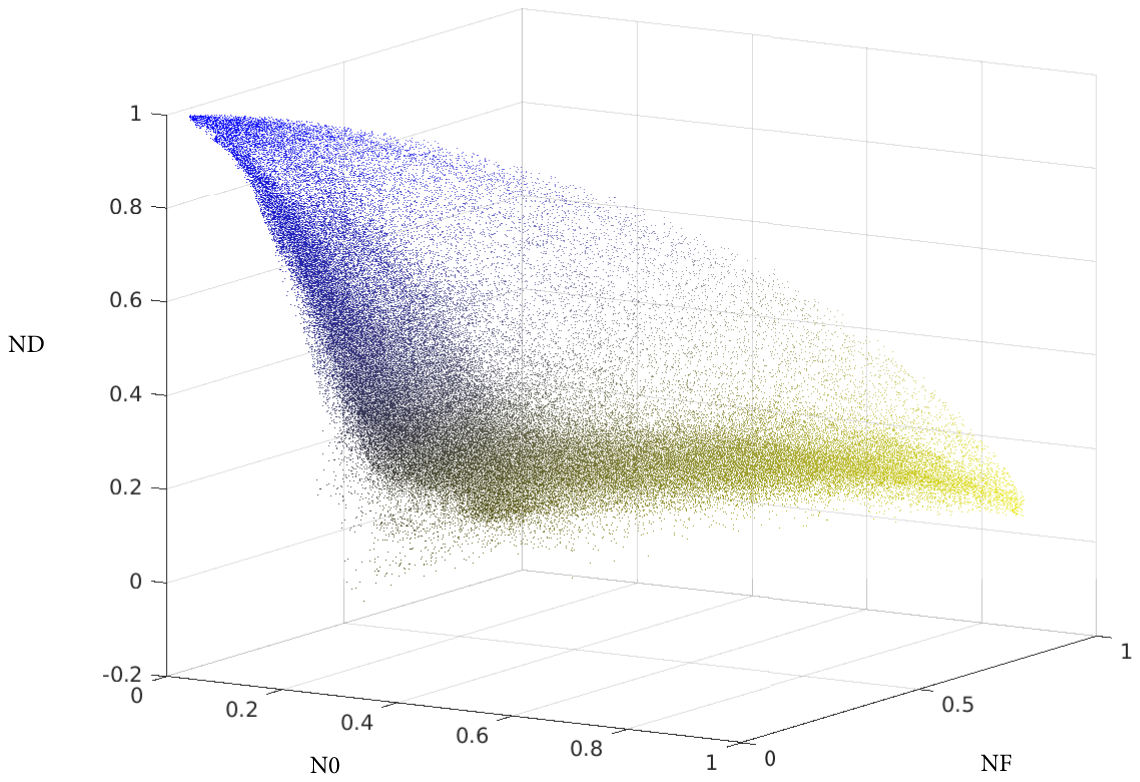

Supplemental Figure 6

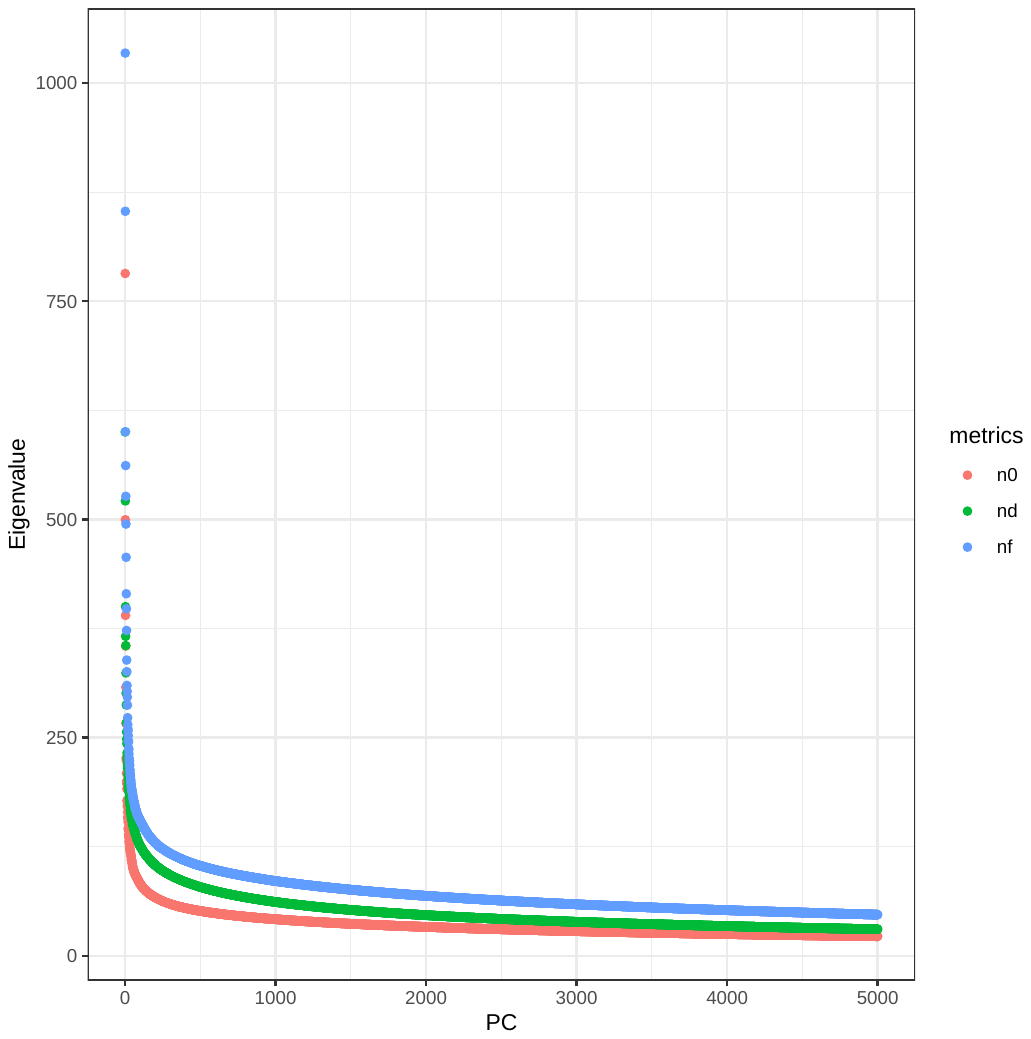

Supplemental Figure 7

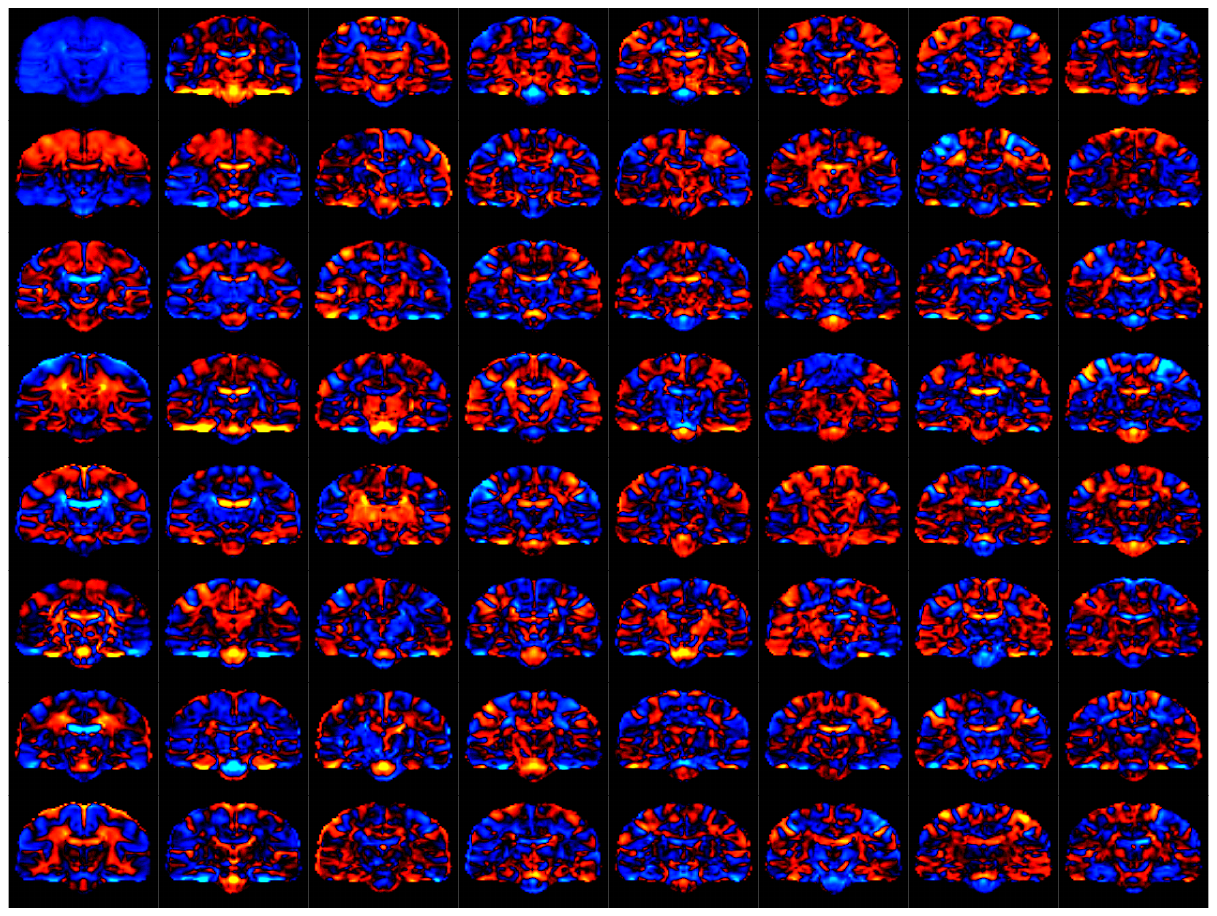

Supplemental Figure 8

Supplemental Figure 9

Supplementary Figure 10

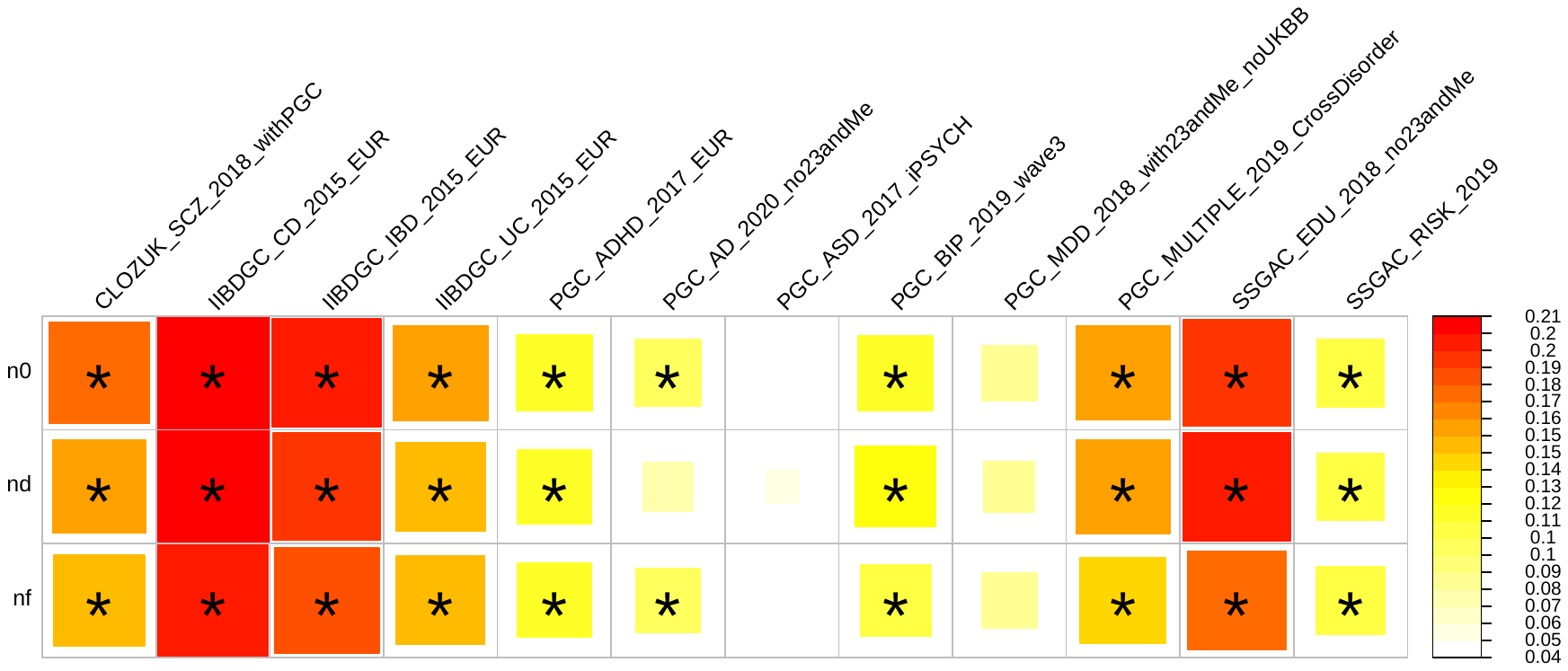

Supplementary Figure 11

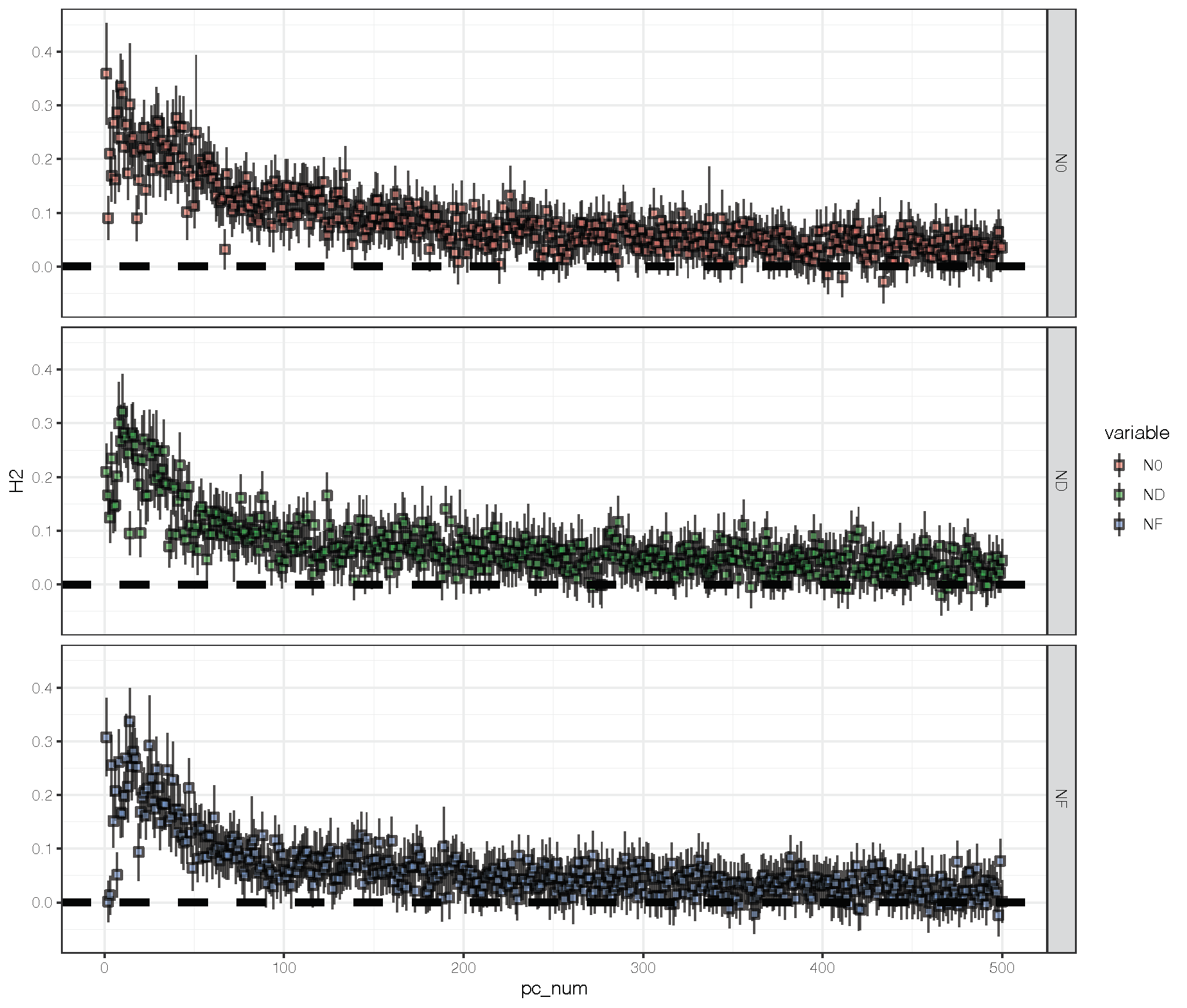
